# Monitoring promoter activity by RNA editing based reporter

**DOI:** 10.1101/2022.03.30.486490

**Authors:** Jing Wang, Yu-Ting Zhao, Lu-Feng Hu, Yangming Wang

**Affiliations:** Institute of Molecular Medicine, College of Future Technology, Peking University, Beijing, China; Academy for Advanced Interdisciplinary Studies, Peking University, Beijing, China

## Abstract

Traditional methods monitoring the promoter activity require the insertion of reporter protein (e.g. fluorescent protein) downstream of a targeted promoter. These approaches suffer from low sensitivity and potential interference to endogenous transcripts especially when the targeted transcript is a noncoding RNA. Here, we develop a mechanistically different reporter system to monitor promoter activity based on ribozyme processed ADAR <u>e</u>ngaging RNA <u>d</u>irected e<u>dit</u>ing (REDDIT). We show that REDDIT can be used to monitor the promoter activity of protein coding, long noncoding RNA, and microRNA (miRNA) genes. Furthermore, REDDIT reporter can also be adapted to use bioluminescence imaging to monitor the promoter activity, which is more suitable for in vivo live imaging. Finally, the reporter sensitivity may be further increased by the circularization of ADAR recruiting RNA. REDDIT provides a powerful platform that is promising in a variety of applications such as monitoring the promoter activity, cell lineage tracing, and processing and manipulating genetic information for synthetic biology applications.

## Introduction

Processing and manipulation of genetic information in living cells is useful for a variety of applications such as cell lineage tracing and genetic circuit engineering^1^. Reporters based on promoter^2^ or miRNA activities^3,4^ have been used to track or manipulate specific cell types. Traditional methods monitor the promoter activity by inserting reporter proteins (e.g. a fluorescent protein) downstream of a specific promoter. These approaches suffer from low sensitivity and potential interference to endogenous transcripts, especially when the targeted transcript is a noncoding RNA. To overcome these problems, we and others recently developed sensitive CRISPR-activator based reporters for endogenous promoters^5,6^. A significant drawback for CRISPR-activator based reporters is that at least three elements have to be introduced, including dCas9-transcriptional activators under the control of ubiquitous strong promoter, a fluorescent protein under the control of an inducible promoter, and a sgRNA precursor inserted downstream of the endogenous promoter. These elements total up to > 9 kb. Here, we develop a mechanistically different reporter system to monitor promoter activity based on ribozyme processed ADAR <u>e</u>ngaging RNA <u>d</u>irected e<u>dit</u>ing (REDDIT). We show that REDDIT can be used to monitor the promoter activity of protein coding, long noncoding RNA and microRNA (miRNA) genes.

## Results

The REDDIT system contains two key elements (**Fig. 1a**): a reporter harboring mCherry and EGFP genes separated by an in-frame UAG stop codon under the control of a constitutively active CAG promoter, and an <u>A</u>DAR <u>R</u>ecruiting-<u>T</u>rigger RNA (ART RNA) that is partially reverse complementary to mCherry-UAG-EGFP reporter. As demonstrated by *Qu* et al^7^, the ART RNA can facilitate the editing of UAG to UIG by recruiting endogenous ADAR, in turn activating the expression of EGFP protein. Theoretically, when the ART RNA is under the control of endogenous promoters, the expression of EGFP should reflect the transcriptional activity of these promoters. We first expressed ART RNA transgenically under the control of a strong CAG promoter. Unfortunately, the editing efficiency by RNA polymerase II transcribed ART RNA was significantly lower than by RNA polymerase III transcribed ART RNA under the control of a U6 promoter (**Fig. S1a-c**). We reasoned that 5’ cap and 3’ polyA structures of pol II transcript may interfere with the editing process. To remove these structures, we inserted self-cleaving twister ribozyme^8^ or tRNA sequences^9^ at both sides of ART RNA (**Fig. 1a**). Although the tRNA flanked ART RNA did not improve the editing efficiency, the editing efficiency by CAG-driven-twister-ribozyme-flanked ART RNA was significantly higher than by either CAG-driven ART RNA or U6-driven ART RNA (**Fig. 1b, c**). Sanger sequencing shows that the editing rate was 17%, 8%, 10%, and 52% for U6-driven, CAG-driven, CAG-driven-tRNA-flanked, and CAG-driven-twister-ribozyme-flanked ART RNAs (**Fig. 1d**), respectively. Moreover, we found that the cleavage by twister ribozymes at both sides are required for high editing activity, since mutations inactivating ribozyme at either side or both sides lowered the editing efficiency to the similar level by CAG-driven ART RNA (**Fig. S2a**).

**Figure 1.**
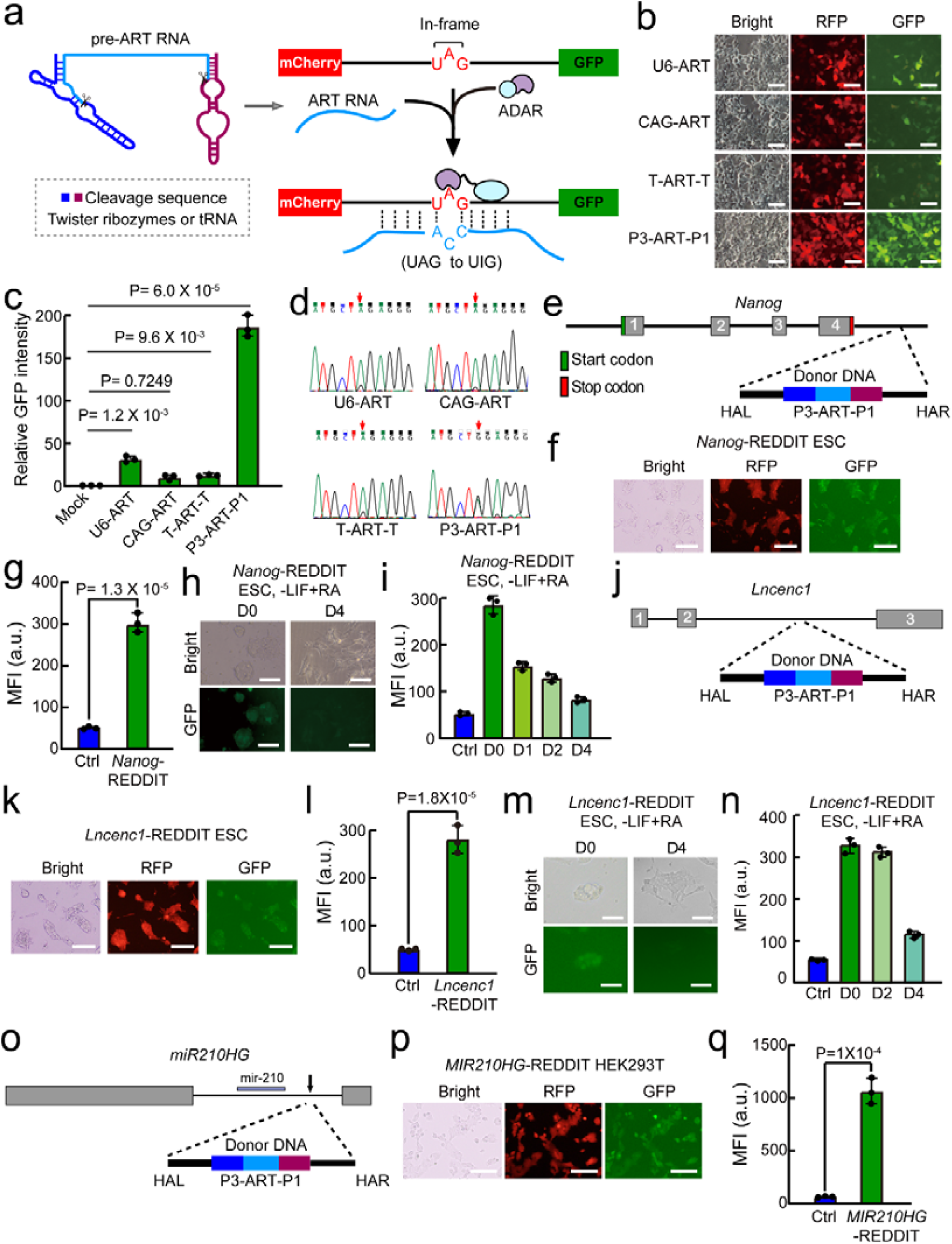
Monitoring the promoter activity of protein coding, lncRNA and miRNA genes by RNA editing based reporter. **a)** Schematic design of REDDIT reporter system. The reporter encode *mCherry* and *GFP* genes separated by an in-frame UAG stop codon. Mature ART RNAs are generated by adding self-cleaving twister ribozyme or tRNA sequences at both sides. mCherry is constitutively expressed, while GFP is not expressed only when the UAG is edited to UIG with the help of ART RNA recruiting endogenous ADAR. Twister ribozyme sequences and tRNA sequences are listed in Table S1. **b)** Representative microscopy images showing the expression of mCherry and GFP in HEK293T transfected with different ART RNA expressing constructs. Scale bars, 200μm. The experiments were repeated three times independently with similar results. U6-ART, U6 promoter driven ART RNA; CAG-ART, CAG promoter driven ART RNA; T-ART-T, CAG promoter driven tRNA flanked ART RNA; P3-ART-P1, CAG promoter driven P3 and P1 twister ribozyme flanked ART RNA. **c)** Quantification of mean GFP intensity of **b**. Shown are mean ± SD, n = 3 independent experiments. The p-value was calculated by one-way ANOVA with two-tailed Dunnett’s test. For mock, ART RNA empty vector was transfected with mCherry-UAG-GFP reporter. **d)** The electropherograms showing Sanger sequencing results in reporter region flanking the UAG codon. The red arrow indicates the edited adenosine of UAG codon. **e)** Schematic of REDDIT knock-in strategy for *Nanog*. After the establishment of transgenic mouse ESCs expressing mCherry-UAG-GFP reporter (serving as control below for *Nanog*-REDDIT ESCs), the twister flanked ART RNA cassette is knocked into *Nanog* locus through CRISPR-Cas9-assisted homologous recombination. **f)** Representative images showing GFP expression in *Nanog*-REDDIT ESCs. Scale bar, 200□μm. **g)** Mean GFP intensity of control and *Nanog*-REDDIT ESCs. The control ESCs express mCherry-UAG-GFP reporter without the knockin of twister ribozyme flanked ART RNA cassette. Shown are mean ± SD, n = 3 independent experiments. The p-value was calculated using two-tailed unpaired Student’s *t* test. **h)** Representative images showing GFP expression in undifferentiated and differentiated *Nanog*-REDDIT mESCs. Scale bar, 200Cμm. **i)** Quantification of mean GFP intensity during the continuous differentiation process of *Nanog*-REDDIT mESCs. Shown are mean ± SD, n = 3 independent experiments. **j)** Schematic of REDDIT knock-in strategy for *Lncenc1*. After the establishment of transgenic mouse ESCs expressing mCherry-UAG-GFP reporter (serving as control below for *Lncenc1*-REDDIT ESCs), the twister flanked ART RNA cassette is knocked in the second intron of *Lncenc1* locus through CRISPR-cas9-assisted homologous recombination. **k)** Representative images showing GFP expression in Lncenc1-REDDIT ESCs. Scale bar, 200□μm. **l)** Mean GFP intensity of control and *Lncenc1*-REDDIT ESCs. The control ESCs express mCherry-UAG-GFP reporter without the knockin of twister ribozyme flanked ART RNA cassette. Shown are mean ± SD, n = 3 independent experiments. The p-value was calculated using two-tailed unpaired Student’s *t* test. **m)** Representative images showing GFP expression in undifferentiated and differentiated *Lncenc1*-REDDIT mESCs. Scale bar, 200□μm. **n)** Quantification of mean GFP intensity during the continuous differentiation process of *Lncenc1*-REDDIT mESCs. Shown are mean ± SD, n = 3 independent experiments. **o)** Schematic of REDDIT knock-in strategy for *MIR210HG*. After the establishment of transgenic HEK293T cells expressing mCherry-UAG-GFP reporter (serving as control for *MIR210HG*-REDDIT HEK293T), the twister ribozyme flanked ART RNA cassette is knocked into *MIR210HG* locus. **p)** Representative images showing GFP expression in *MIR210HG*-REDDIT HEK293T cells. Scale bar, 200□μm. **q)** Mean GFP intensity of control and *MIR210HG*-REDDIT HEK293T cells. The control HEK293T cells express mCherry-UAG-GFP reporter without the knockin of twister ribozyme flanked ART RNA cassette. Shown are mean ± SD, n = 3 independent experiments. The p-value was calculated using two-tailed unpaired Student’s *t* test.

Next, we evaluated the requirement for the length of ART RNA. The ART RNA used above is 161 nt long, which is optimal for editing based on previous study by Qu et al^7^. Consistent with this study, longer ART RNA induces higher editing efficiency (**Fig. S2b**). Based on these data, the 161 nt ART RNA was chosen for further experiments in this study. We then tested whether twister ribozyme flanked ART RNA can also trigger the editing of endogenous genes. For two sites on PPIB and KRAS, comparable editing efficiency was achieved by U6-driven and CAG-driven twister ribozyme flanked ART RNA in HEK293T cells (**Fig. S3a, b**). Finally, when the twister ribozyme flanked ART RNA was placed under the control of a tetracycline inducible promoter, the expression of GFP reporter is significantly induced upon the addition of doxycycline (**Fig. S4a-c**). More importantly, the EGFP expression correlated with the dosage of doxycycline (**Fig. S4d**), supporting that REDDIT can be used to monitor the promoter activity.

Next, we tested whether REDDIT can be used to monitor the activity of endogenous promoters. We first knocked the twister ribozyme-flanked ART RNA at the 3’ untranslated region of *Nanog* in mouse embryonic stem cells (ESCs) in which mCherry-UAG-EGFP is transgenically expressed under the control of a CAG promoter (**Fig. 1e**). Nanog is a pluripotency gene enriched in ESCs but downregulated during differentiation^10^. In ESCs with REDDIT successfully integrated at *Nanog* locus (*Nanog*-REDDIT ESCs), we observed high expression level of GFP (**Fig. 1f, g**). Importantly, the knock-in of REDDIT has no interference for the expression of Nanog and other pluripotency genes including Oct4 (also known as Pou5f1) and Sox2 (**Fig. S5a**). When the differentiation was induced by all-trans retinoid acids (ATRA) in the absence of leukemia inhibitory factor (LIF), the Nanog expression was significantly decreased (**Fig. S5b**). Correspondingly, the *Nanog*-REDDIT EGFP level was also decreased upon differentiation (**Fig. 1h, i**). Together, these results suggest that REDDIT can monitor the promoter activity of protein coding genes.

Next, we inserted the twister ribozyme-flanked ART RNA to the second intron of a long noncoding RNA *Lncenc1*^11^ (**Fig. 1j**). Similarly, EGFP was significantly induced in *Lncenc1*-REDDIT ESCs (**Fig. 1k, l**). Again, the insertion of twister ribozyme-flanked ART RNA did not interfere the expression of Lncenc1 or pluripotency genes (**Fig. S6a**). Upon differentiation, Lncenc1 was significantly downregulated in *Lncenc1*-REDDIT ESCs (**Fig. S6b**). Consistently, the EGFP level was also significantly downregulated (**Fig. 1m, n**). In addition, cells with high EGFP level expressed high level of Lncenc1 (**Fig. S6c, d**). Together, these data suggest that REDDIT can monitor the promoter activity of long noncoding RNAs.

Finally, we made a REDDIT reporter for *MIR210HG* gene (**Fig. 1o**), which encodes a miRNA important for cancer progression^12^. Again, EGFP expression was highly induced in *MIR210HG*-REDDIT HEK293T cells (**Fig. 1p, q**). Together, these data show that REDDIT may be used to monitor the promoter activity of miRNA genes.

Bioluminescence imaging has been widely used in vivo for its noninvasive nature. We then tested whether the EGFP can be replaced by luciferase (**Fig. S7a**). A CAG promoter driven twister ribozyme-flanked ART RNA triggered ∼580 fold increases in the expression of luciferase (**Fig. S7a-c**). Moreover, using a previously reported strategy to circularize ART RNA, the sensitivity for REDDIT reporter is further increased by ∼34 fold (**Fig. S7a-c**). These data show that REDDIT reporter may be adapted to use bioluminescence imaging to monitor the promoter activity, which is more suitable for in vivo live imaging.

## Conclusion and Discussion

In summary, REDDIT is a novel strategy to monitor the promoter activity of different category of genes including protein-coding, long noncoding RNA and miRNA genes. Compared to traditional method inserting a fluorescent protein gene, the REDDIT strategy inserts a much smaller fragment (281 nt) into the endogenous DNA locus. Theoretically, REDDIT should cause less interference for the expression or function of noncoding RNAs, since it does not require the noncoding RNA being reprogrammed to a coding gene expressing fluorescent proteins. In addition, REDDIT may be used to monitor the promoter in vivo through non-invasive bioluminescence imaging when EGFP is replaced with luciferase. Finally, if EGFP is replaced with other genes, for example, transcription factors or RNA binding proteins, REDDIT may be used to build synthetic genetic circuit to control gene expression and cell behavior. In conclusion, REDDIT provides a powerful platform that is promising in a variety of applications such as monitoring the promoter activity, cell lineage tracing, processing and manipulating genetic information for synthetic biology applications.

## Acknowledgements

We would like to thank members of Wang laboratory for discussion of the project. We thank the Flow Cytometry Core at National Center for Protein Sciences at Peking University, particularly Huan Yang, Jia Luo, Xuefang Zhang, Yinghua Guo and Dr. Hongxia Lv for technical help. We thank Yiwen Zhang for assistance in constructing plasmids and cell culture. This study was supported by The National Key Research and Development Program of Ministry of Science and Technology of the People’s Republic of China [2021YFA0100200 and 2018YFA0107601] and National Natural Science Foundation of China [91940302, 32130017 and 32025007] to Yangming Wang.

## Author contributions

Jing Wang and Lu-Feng Hu performed experiments in Figures 1c, 1d, S1b, S1c, S2, S3, S4b and S4c. Yu-Ting Zhao performed experiments in Figures 1j-n, S4d, S6 and S7. Jing Wang also performed experiments in Figures 1e-i, 1o-1q, S4a-c and S5. Jing Wang and Yu-Ting Zhao draw all the figures with the help from Lu-Feng Hu. All the authors interpreted the data. Yangming Wang conceived and supervised the project and wrote the manuscript with the help from Jing Wang, Yu-Ting Zhao and Lu-Feng Hu.

## Supplementary Information

**Figure S1.**
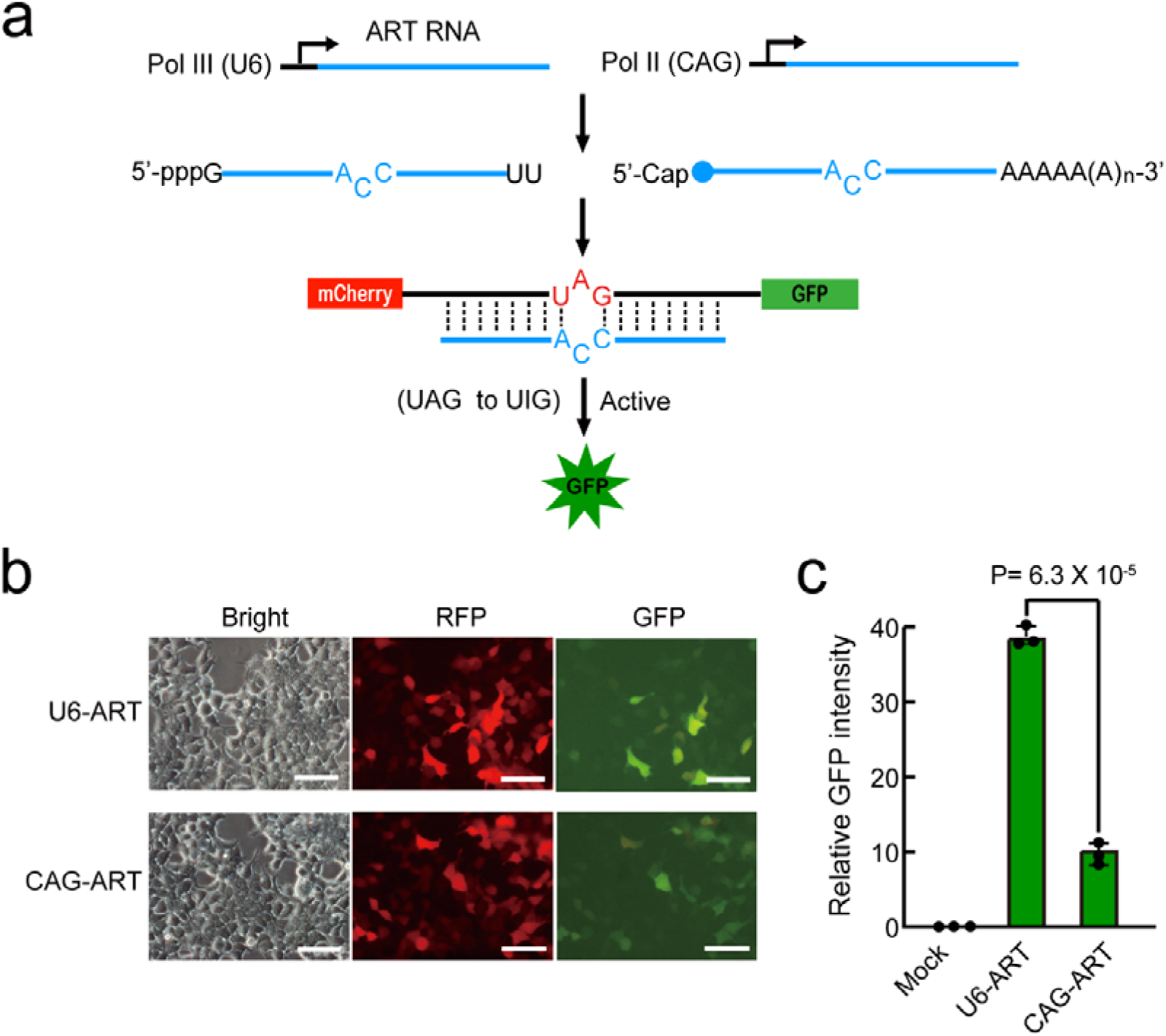
CAG promoter construct is less efficient than U6 promoter construct in triggering RNA editing. **a)** Schematic graph showing ART RNA driven by Pol II or Pol III promoter in triggering the activation of GFP through RNA editing. **b)** Representative microscopy images showing mCherry and GFP expression in HEK293T transfected with mCherry-UAG-GFP reporter and U6 or CAG promoter constructs. Scale bars, 200μm. **c)** Quantification of mean GFP intensity. Shown are mean ± SD, n = 3 independent experiments. The p-value was calculated using two-tailed unpaired Student’s *t*-test.

**Figure S2.**
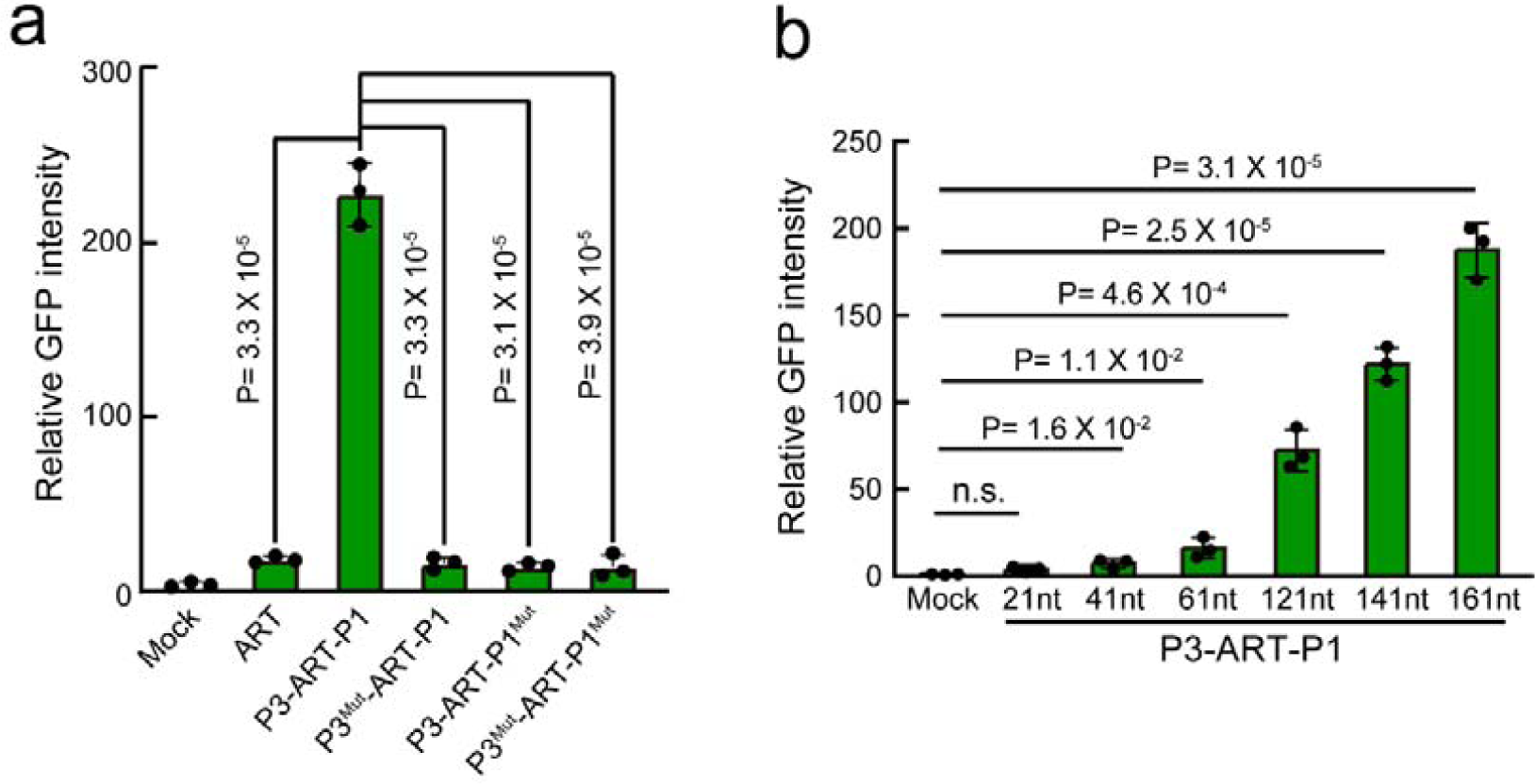
Characterization and optimization of ART RNA expression construct. **a)** Mean GFP intensity showing the editing efficiency by constructs producing variable precursor ART RNAs in HEK29T. For twister flanked ART RNA, mutations are introduced into 5’ or 3’ twister ribozyme to inactivate the clevage activity; Shown are mean ± SD, n = 3 independent experiments. The p-value was calculated by one-way ANOVA with two-tailed Dunnett’s multiple comparison test. **b)** Relative GFP intensity showing the editing efficiency by constructs producing ART RNAs with different length. All ART RNAs are flanked by twister riboyzme at both sides. Data were normalized to HEK293T transfected with mCherry-UAG-GFP and empty ART RNA constructs. The p-value was calculated by one-way ANOVA with two-tailed Dunnett’s multiple comparison test.

**Figure S3.**
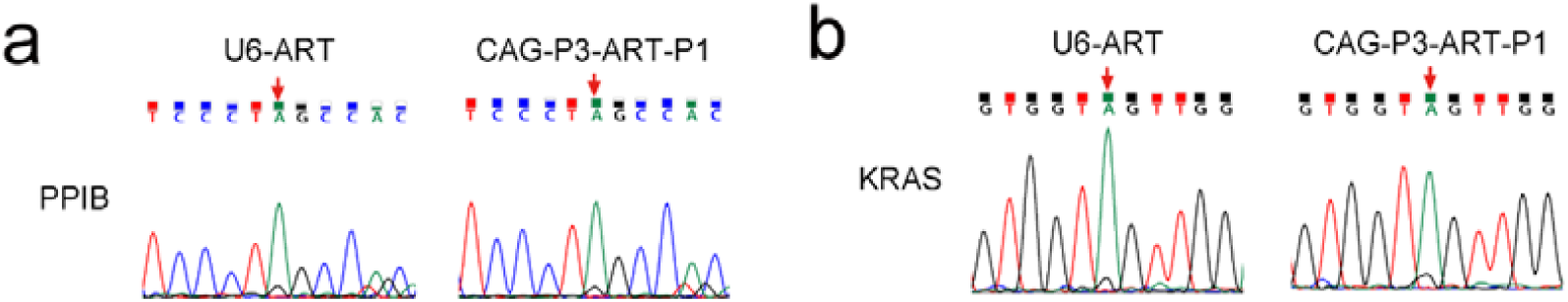
The editing efficiency on edogenous transcripts is similar between CAG promoter driven twister ribozyme flanked ART RNA and U6 promoter driven ART RNA in HEK293T cells. **a)** Sanger sequencing results showing the editing rate on targeted adenosine of the endogenous PPIB transcript by U6 promoter driven ART RNA or CAG promoter driven twister ribozyme flanked ART RNA. The red arrow indicates the targeted adenosine. **b)** Sanger sequencing results showing the editing rate on targeted adenosine of the endogenous KRAS transcript by U6 promoter driven ART RNA or CAG promoter driven twister ribozyme flanked ART RNA. The red arrow indicates the targeted adenosine.

**Figure S4.**
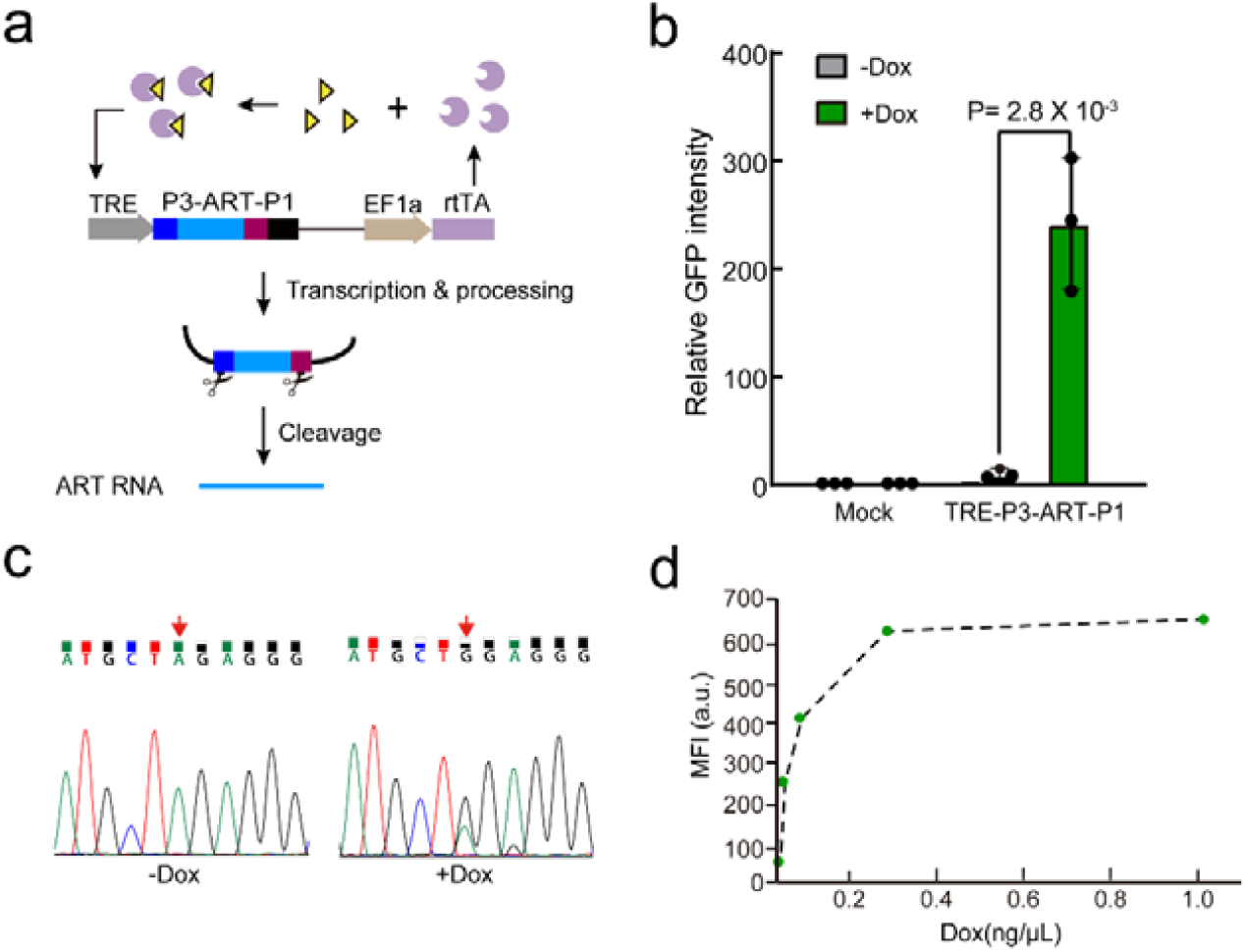
Inducible RNA editing is enabled by producing twister ribozyme flanked ART RNA through a doxycycline inducible promoter. **a)** Schematic representation of tetracycline-inducible system. The ART RNA precursor is placed under the control of an rtTA-responsive promoter TRE3G. In the presence of doxycycline, the rtTA protein driven by EF1a promoter binds to TRE promoter to activate the transcription of precursor ART RNA. **b)** Mean GFP intensity showing inducible RNA editing. The mock control is HEK293T transfected with mCherry-UAG-GFP and empty inducible construct not expressing any ART RNA. Shown are mean ± SD, n = 3 independent experiments. The p-value was calculated by two-tailed unpaired Student’s t-test. **c)** Sanger sequencing results showing the editing rate on targeted adenosine of the reporter transcripts in the absence or presence of doxycycline. Beacuse the editing efficiency is more than 50%, the base A position is recognized as G in the sequencing for Dox+ sample. **d)** Mean fluorescence intensity of GFP at different Dox concentrations 48□h after Dox addition.

**Figure S5.**
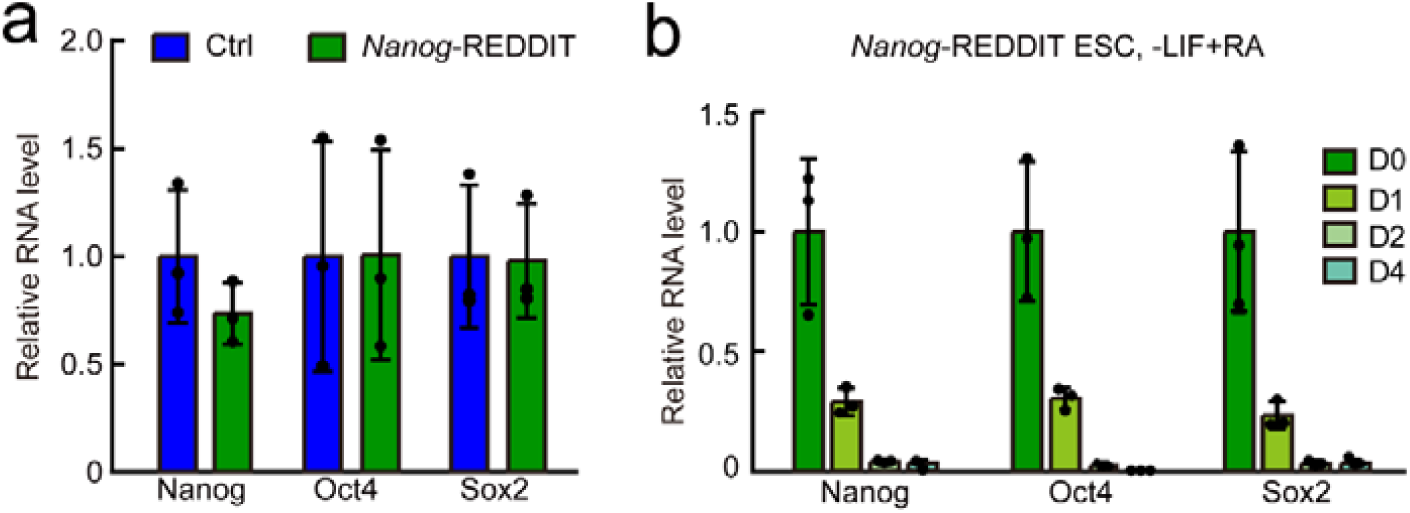
Twister ribozyme flanked ART RNA knocked in *Nanog* locus has little impact on the expression of endogenous Nanog and can track the differentiation status of ESCs. **a)** RT-qPCR analysis of Nanog, Oct4 and Sox2 in control and *Nanog*-REDDIT ESCs. Control ESCs refer to embryonic stem cell line stably expressed mCherry-UAG-GFP reporter with no twister flanked ART RNA knocked in *Nanog* locus. β-actin mRNA was used as a control. Data were normalized to control ESCs. Shown are mean ± SD, n = 3 independent experiments. **b)** RT-qPCR analysis of Nanog, Oct4 and Sox2 during differentiation process of *Nanog*-REDDIT ESCs. Shown are mean ± SD, n = 3 independent experiments.

**Figure S6.**
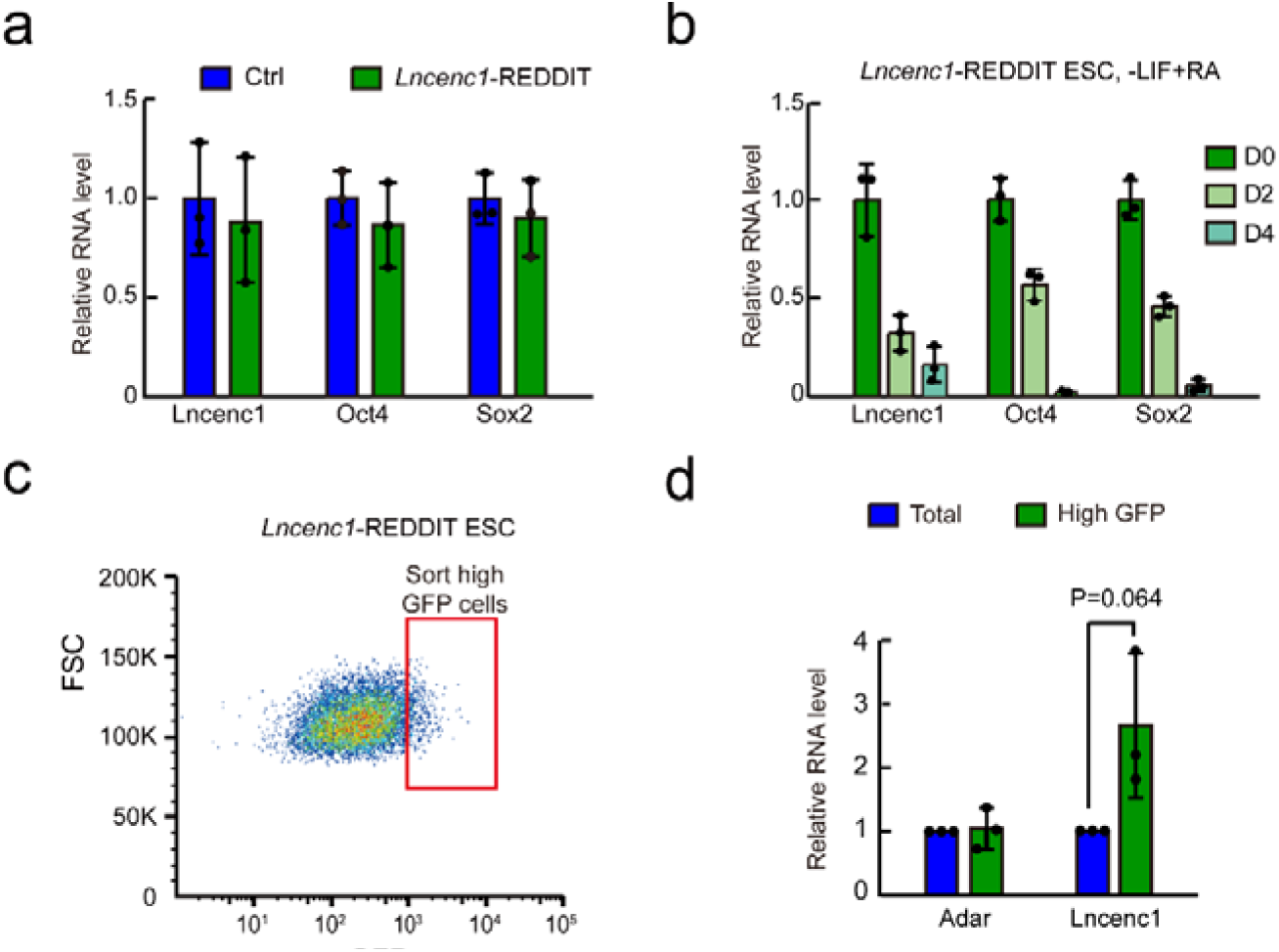
Twister ribozyme flanked ART RNA knocked in *Lncenc1* locus has little impact on the expression of endogenous Lncenc1 and can track the differentiation status of ESCs. **a)** RT-qPCR analysis of Lncenc1, Oct4 and Sox2 in control and *Lncenc1*-REDDIT ESCs. Control ESCs refer to embryonic stem cell line stably expressed mCherry-UAG-GFP reporter with no twister ribozyme flanked ART RNA knocked in *Lncenc1* locus. β-actin mRNA was used as a control. Data were normalized to control ESCs. Shown are mean ± SD, n = 3 independent experiments. **b)** RT-qPCR analysis of Lncenc1, Oct4 and Sox2 during differentiation process of *Lncenc1*-REDDIT ESCs. Shown are mean ± SD, n = 3 independent experiments. **c)** Representative flow cytometry scatter plot and sorting gate of GFP high cells for *Lncenc1*-REDDIT ESCs. **d)** RT-qPCR analysis of Lncenc1 in GFP high or total population of *Lncenc1*-REDDIT ESCs. Data were normalized to β-actin and then to total population. Shown are mean ± SD, n = 3 independent experiments. The p-value was calculated using two-tailed paired Student’s *t* test.

**Figure S7.**
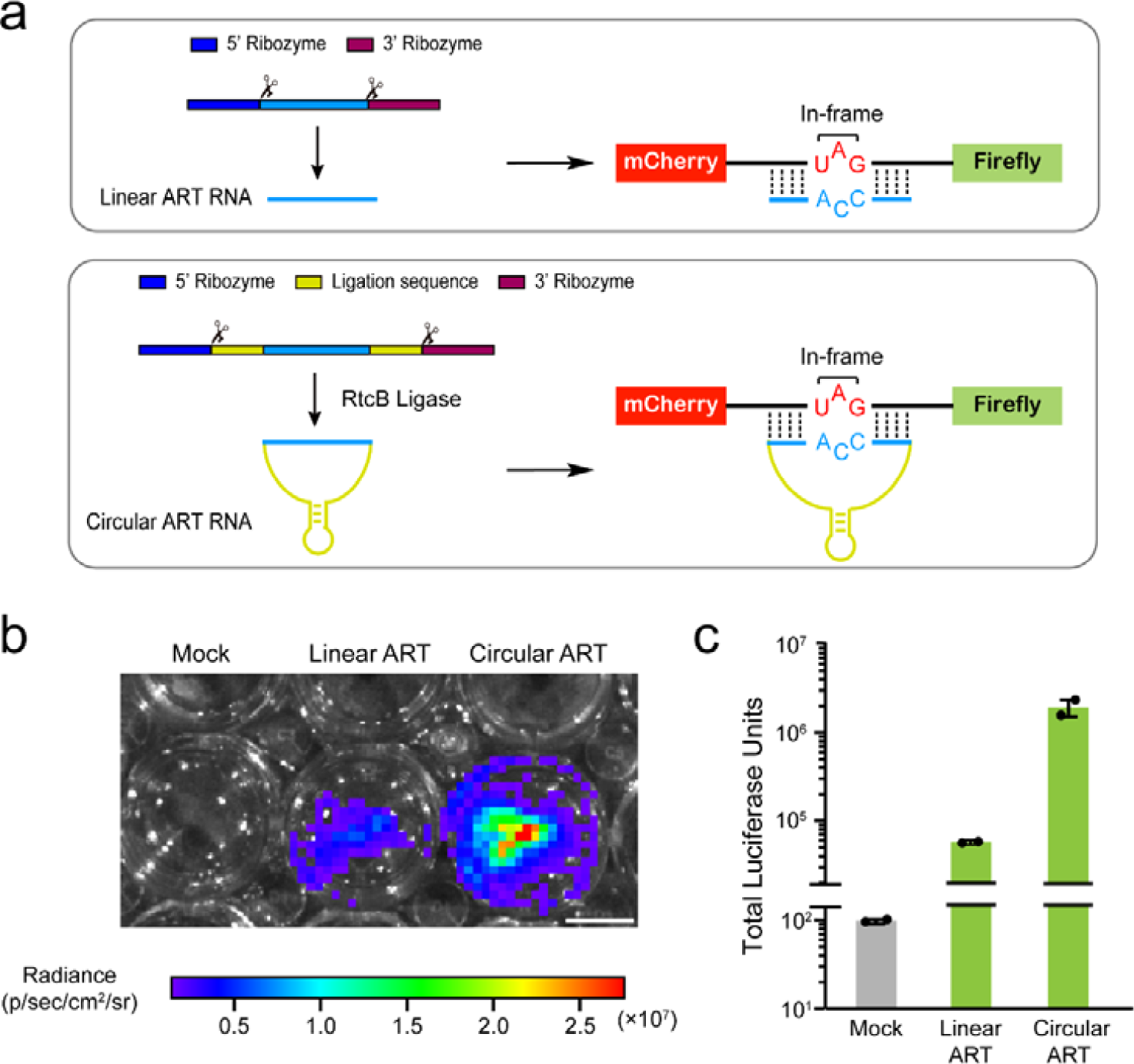
REDDIT can be adapted to bioilluminescence imaging. **a)** Schematic representation of the luciferase expression activated by linear or circular ART RNA. The precursor of circular ART RNA is produced by CAG pol II promoter and processed by twister flanked ribozyme. After processing, the synthesis of circular RNA is achieved through the ligation of 5’ and 3’ end by the endogenous RNA ligase RtcB. **b)** Representative luminescence images showing the activation of firefly luciferase by linear or circular ART RNA in HEK293T. For mock, only mCherry-UAG-Firefly reporter was transfected into cells. Scale bar, 1 cm. **c)** Quantification of mean luciferase activity. n = 2 independent experiments.

### Supplementary Materials and Methods

#### Plasmid and vector construction

The mCherry-TAG-GFP reporter sequence is the same as reported previously and was cloned downstream of the CAG promoter in a piggyBac vector containing a Zeocin resistance gene. The ADAR recruiting-trigger RNA (ART RNA) and tRNA or twister ribozyme flanked ART RNAs, either targeting mCherry-TAG-GFP reporter or endogenous RNAs, were cloned downstream of the CAG promoter or U6 promoter as indicated in Figures. The inducible ART RNA targeting mCherry-TAG-GFP reporter was driven by the doxycycline (Dox) responsive TRE3G promoter in a piggyBac vector containing a hygromycin resistance gene. To construct circular ART RNA, a similar strategy as reported by Litke et al. was adopted, in which ART RNA is flanked by the 5’- and 3’-stem-forming sequences each of which was flanked by a twister ribozyme.

To construct donor template for knocking ART RNA precursor into endogenous gene locus with CRISPR-Cas9, the 5’ and 3’ homology arms were PCR amplified from mouse or human genomic DNA. For *Nanog*-REDDIT, restriction endonuclease sites PacI and SalI were introduced into the middle of homology arm by PCR. Then the homology arms with restriction endonuclease sites were cloned into the pCE2 vector with TA/Blunt-Zero Cloning Kit (Vazyme, Cat. #C601). Meanwhile, the PCR products of ART RNA were digested with PacI and SalI (NEB, Cat. # R0547 and Cat. # R0560) and purified. The PacI and SalI digested vector with homology arms and ART RNA PCR products were then ligated using T4 ligase (Thermofisher, Cat. # EL0012) to construct *Nanog*-REDDIT knock in template. For *Lncenc1*-REDDIT, the procedure for constructing the donor template was essentially the same as *Nanog*-REDDIT except that restriction endonuclease sites PacI and PmeI instead of PacI and SalI were used. For *MIR210HG*-REDDIT, the procedure for constructing donor template was same as that of *Nanog*-REDDIT. The sgRNAs for Cas9-mediated knock-in were designed by CHOPCHOP (https://chopchop.cbu.uib.no/) and were cloned into pXG2 vector. All the PCR products were generated by Phanta® HS Super-Fidelity DNA Polymerase (Vazyme, Cat. #P502-d) and purified by a FastPure Gel DNA Extraction (Vazyme, Cat. #DC301).

#### Cell culture and construct of reporter cell lines

HEK293T cells were cultured at 37 °C under 5% CO_2_ in high glucose Dulbecco’s Modified Eagle’s Medium (DMEM, Hyclone, Cat. #SH30243.01) supplemented with 10% FBS (PANSera, Cat. #2602-P130707). Embryonic stem cells were grown on gelatin-coated dish at 37 °C under 5% CO2, in KnockOut™ DMEM (Gibco, Cat. # 10829081) supplemented with 15% FBS (Hyclone, Cat. # SH3007103), 1,000 U/ml mouse leukemia inhibitory factor (1,000 U/ml), 0.1 mM non-essential amino acids (Gibco, Cat. # 11140050), 1 mM L-glutamine (Gibco, Cat. # 25030081), 0.1 mM β-mercaptoethanol, and penicillin (100 U/ml) and streptomycin (100 µg/ml).

To establish REDDIT reporter ESCs, we first generated mouse V6.5 ESCs in which mCherry-UAG-GFP was stably integrated. The mCherry-UAG-GFP plasmid was transfected with PBase expression plasmid using Polyplus Transfection (Jetprime, Cat. # PT-114-75) reagent. After transfection, cells were treated with 20 µg/ml Zeocin (Beyotime, Cat. # ST1450) for 5 days. Then monoclonal cell line was selected and expanded for further experiments. The REDDIT cassettes were then knocked in the host gene locus assisted with CRISPR/*Spy*Cas9 system through HDR (homology-directed repair) strategy. For *Nanog*-REDDIT, ART RNA cassette was inserted downstream of poly (A). For *Lncenc1*-REDDIT ESCs, ART RNA cassette was knocked into the second intron of Lncenc1. Establishment procedure of *MIR210HG*-REDDIT HEK293T reporter cell line was similar as that of REDDIT reporter ESCs. ART RNA cassette was knocked in around 500 bases downstream of pre-miR-210 sequence.

For monolayer retinoid acid (RA) differentiation, 50,000 ESCs per well were seeded in ESC media in 6-well plates. After 24h, the medium was changed to differentiation media without LIF and in the presence of 100 nM all-trans retinoid acid. *Lncenc1*-REDDIT ESCs and *Nanog*-REDDIT ESCs were differentiated continuously for 4 days. Medium was changed every day.

#### RNA editing of reporter or endogenous RNA

To edit reporter transcripts, 50,000 HEK293T cells were plated on poly-D-lysine (PDL) coated 24-well. 50ng mCherry-TAG-GFP reporter plasmid and 450ng ART RNA plasmid were co-transfected with Polyplus Transfection (Jetprime, Cat. # PT-114-75). At 48h post transfection, cells were harvested and then analyzed with flow cytometry and Sanger sequencing. For Dox inducible RNA editing, 250,000 HEK293T cells were seeded in 12-well plate before transfection. After 16h, 100 ng mCherry-TAG-GFP reporter plasmid and 400 ng TRE-ART RNA plasmid were co-transfected by Polyplus Transfection. 24h after transfection, fresh media contain 20 µg/ml Zeocin was added to enrich the mCherry-TAG-GFP positive cells. Two days later, cells were digested and plated in fresh medium containing doxycycline at indicated concentrations. 48h after Dox treatment, cells were harvested and analyzed with flow cytometry.

To edit endogenous transcripts, 50,000 HEK293T cells were plated on poly-D-lysine (PDL) coated 24-well. 50ng mCherry-TAG-GFP reporter plasmid and 450ng ART RNA plasmid were co-transfected with Polyplus Transfection (Jetprime, Cat. # PT-114-75). At 48h post transfection, cells were harvested and RNA extraction was conducted. To evaluate RNA editing efficiency, the total RNA were reverse transcribed into cDNA and then the region containing the targeted editing site was PCR amplified. The PCR products were purified by agarose electrophoresis before submitted for Sanger sequencing (Azenta Life Sciences).

#### RNA extraction and RT-qPCR

Total RNA was extracted from cells with Trizol reagent (Magen, Cat. # R4801). For quantitative PCR with reverse transcription (RT-qPCR) analysis, 250-500 ng total RNA were reverse transcribed using HiScript III RT SuperMix for qPCR (Vazyme, Cat. # R323). qPCR assay was carried out with AceQ qPCR SYBR Green Mater Mix reagent (Vazyme, Cat. # Q141) in 96-well plates on StepOne Plus Real-Time PCR System (Applied Biosystems) with standard protocols.

#### Fluorescence activated cell sorting and flow cytometry analysis

Cells were first dissociated with 0.1% Trypsin and then were washed twice with PBS. The cells were resuspended in PBS containing 2% FBS. Flow cytometry for quantifying population and fluorescence intensity was performed by BD LSR Fortessa SORP. Cell sorting was performed on BD FACS Aria III. Data were analyzed using FlowJo software.

#### Bioluminescence imaging and luciferase assay system

Before transfection, 300,000 HEK293T cells per well were seeded in 12-well plates. After 16h, the linear ART RNA or circular ART RNA plasmids were co-transfected along with mCherry-UAG-Luciferase encoding plasmid using Polyplus Transfection reagent. The transfected cells were enriched with 130μg/ml Hygromycin (Roche) for 2 days. Bioluminescence imaging was performed with IVIS® Optical Imaging System (Perkin Elmer). Before imaging, cells were incubated in fresh medium containing 150μg/ml D-Luciferin potassium salt (Solarbio Cat. # D8390) for 5min at room temperature. For measuring luciferase activity, 20,000 HEK293T cells were processed with reagents from Dual Luciferase Reporter Assay Kit (Promega, Cat. # E1531) and measured with DLReady Luminometer (Berthold).

#### Quantification and statistical analysis

The data were presented as mean ± SD, except where indicated otherwise. Statistical analyses were performed using the GraphPad Prism v6 software. Statistical significance was assessed by two-tailed Student’s *t*-test. For multiple comparison, the p-value was calculated by one-way ANOVA followed with Dunnett’s test or Tukey’s multiple comparison test. p<0.05 is generally considered as statistically significant. FlowJo_V10 software (BD Biosciences)was used for flow cytometry analysis.

#### Data and materials availability

All data generated or analyzed during this study are included in the manuscript and its supplementary information files. Original data used and/or analyzed during the current study are available from the corresponding author on reasonable request.

## References

1 Black, J. B., Perez-Pinera, P. & Gersbach, C. A. Mammalian Synthetic Biology: Engineering Biological Systems. Annu Rev Biomed Eng 19, 249–277, doi:10.1146/annurev-bioeng-071516-044649 (2017).

2 Liu, H. et al. Systematically labeling developmental stage-specific genes for the study of pancreatic beta-cell differentiation from human embryonic stem cells. Cell Res 24, 1181–1200, doi:10.1038/cr.2014.118 (2014).

3 Wang, X. W. et al. A microRNA-inducible CRISPR-Cas9 platform serves as a microRNA sensor and cell-type-specific genome regulation tool. Nat Cell Biol 21, 522–530, doi:10.1038/s41556-019-0292-7 (2019).

4 Miki, K. et al. Efficient Detection and Purification of Cell Populations Using Synthetic MicroRNA Switches. Cell Stem Cell 16, 699–711, doi:10.1016/j.stem.2015.04.005 (2015).

5 Gao, N. et al. Endogenous promoter-driven sgRNA for monitoring the expression of low-abundance transcripts and lncRNAs. Nat Cell Biol 23, 99–108, doi:10.1038/s41556-020-00610-9 (2021).

6 Zhao, Y. T. & Wang, Y. Monitoring the promoter activity of long noncoding RNAs and stem cell differentiation through knock-in of sgRNA flanked by tRNA in an intron. Cell Discov 7, 45, doi:10.1038/s41421-021-00272-3 (2021).

7 Qu, L. et al. Programmable RNA editing by recruiting endogenous ADAR using engineered RNAs. Nat Biotechnol 37, 1059–1069, doi:10.1038/s41587-019-0178-z (2019).

8 Roth, A. et al. A widespread self-cleaving ribozyme class is revealed by bioinformatics. Nat Chem Biol 10, 56–60, doi:10.1038/nchembio.1386 (2014).

9 He, X. et al. Boosting activity of high-fidelity CRISPR/Cas9 variants using a tRNA(Gln)-processing system in human cells. J Biol Chem 294, 9308–9315, doi:10.1074/jbc.RA119.007791 (2019).

10 Loh, Y. H. et al. The Oct4 and Nanog transcription network regulates pluripotency in mouse embryonic stem cells. Nat Genet 38, 431–440, doi:10.1038/ng1760 (2006).

11 Sun, Z. et al. The Long Noncoding RNA Lncenc1 Maintains Naive States of Mouse ESCs by Promoting the Glycolysis Pathway. Stem cell reports 11, 741–755, doi:10.1016/j.stemcr.2018.08.001 (2018).

12 Dang, K. & Myers, K. A. The role of hypoxia-induced miR-210 in cancer progression. Int J Mol Sci 16, 6353–6372, doi:10.3390/ijms16036353 (2015).

